# Cluster Headache: Comparing Clustering Tools for 10X Single Cell Sequencing Data

**DOI:** 10.1101/203752

**Authors:** Saskia Freytag, Ingrid Lonnstedt, Milica Ng, Melanie Bahlo

## Abstract

The commercially available 10X Genomics protocol to generate droplet-based single cell RNA-seq (scRNA-seq) data is enjoying growing popularity among researchers. Fundamental to the analysis of such scRNA-seq data is the ability to cluster similar or same cells into non-overlapping groups. Many competing methods have been proposed for this task, but there is currently little guidance with regards to which method offers most accuracy. Answering this question is complicated by the fact that 10X Genomics data lack cell labels that would allow a direct performance evaluation. Thus in this review, we focused on comparing clustering solutions of a dozen methods for three datasets on human peripheral mononuclear cells generated with the 10X Genomics technology. While clustering solutions appeared robust, we found that solutions produced by different methods have little in common with each other. They also failed to replicate cell type assignment generated with supervised labeling approaches. Furthermore, we demonstrate that all clustering methods tested clustered cells to a large degree according to the amount of genes coding for ribosomal protein genes in each cell.

## Introduction

Single cell RNA-sequencing (scRNA-seq) technology theoretically allows us to comprehensively catalog every cell type on the planet (1) and, in the future, from other planets. Indeed, one effort underway, the Human Cell Atlas, aims to characterize molecular composition and origin of every cell type within the human body (2), a daunting goal in itself. At the heart of such endeavors is the ability to identify both known and novel cell types. This is accomplished using computational methods that characterize cells in heterogeneous tissue samples examined by scRNA-seq. Characterization consists of two steps: (i) clustering of same or similar cells into non-overlapping groups, and (ii) labeling clusters, i.e. determining the cell type represented by the cluster. Here, we focus on the first step of this process.

The vast majority of computational methods for scRNA-seq cell type clustering can be categorized as either unsupervised or semi-supervised clustering strategies (3). Unsupervised methods do not use any prior information to cluster cell into groups. In other words they use the whole transcriptome agnostically to separate cells of different types. In contrast, semi-supervised methods make use of existing information about cell types. Some semi-supervised methods use the transcriptome in a more discriminate manner, for example by exclusively focusing on subsets of genes differentially expressed in different cell types. Other semi-supervised methods project data into a space defined by expression data on isolated cell types and then apply clustering. Note that fully supervised methods are not generally suited for the task of clustering scRNA-seq data as they require knowledge on all cell types present within the sample. In particular, use of supervised methods precludes the discovery of new cell types.

Research into clustering has produced many algorithms for the task. These algorithms vary considerably in complexity and with regards to their assumptions. There are few rules guiding the application of clustering algorithms to a particular problem, such as clustering of scRNA-seq. This combined with the relative youth of the field mean no consensus as to which clustering method is most accurate has been reached thus far. Furthermore, the existence of a universally best clustering algorithm for scRNA-seq is doubtful, as different scRNA-seq protocols differ in terms of throughput capabilities and sensitivities (4). It is likely that catering for the intricacies of a particular protocol can always improve performance of clustering on data generated by said protocol. Indeed, most clustering methods are developed and tested on only one scRNA-seq protocol.

Here we focus solely on data generated by a protocol commercialized by 10X Genomics. Commercially available scRNA-seq platforms, like 10X Genomics’ Chromium, are being widely adopted due to their ease of use and relatively low cost per cell (5). The 10X Genomics protocol uses a droplet-based system to isolate single cells. Each droplet contains all the necessary reagents for cell lysis, barcoding, reverse transcription and molecular tagging. This is followed by pooled PCR amplification and 3’ library preparation, after which standard Illumina short-read sequencing can be applied (6). Unlike other commercially available scRNA-seq protocols, like Fluidigm C1, 10X Genomics allows sequencing of thousands of cells albeit at much shallower read depth per cell. As such the 10X platform is particularly suited to detailed characterization of heterogeneous tissues.

## Methods and Materials

### Datasets and Preprocessing

We analyzed three scRNA-seq datasets, summarized in Table 1, examining human peripheral mononuclear blood cells (PBMCs) from healthy donors. All datasets were generated using the 10X Genomics droplet system combined with Illumina sequencing. The first two datasets were generated by 10X Genomics and are publicly available. Of these one dataset was generated with an earlier version of the microfluidics instrument. This dataset will be referred to as the 10X GemCode dataset. The second dataset was generated with the latest instrument, the 10X Chromium (thus the data will be referred to as 10X Chromium). The Australian Genome Research Facility in partnership with CSL generated the third dataset using the 10X Chromium system. This dataset will be referred to as CSL data from now on. We also had access to a bulk RNA-seq dataset containing the expression of 8 isolated cell types found in PBMCs [de Graaf et al, Stem Cell Reports 2016]. Datasets containing the expression of isolated cell types can serve as references for the analysis of scRNA-seq data.

**Table 1.** Comparison of three scRNA-seq datasets investigating PBMCs from healthy donors

| Dataset Name | CSL Data | 10X GemCode Data | 10X Chromium Data |
| --- | --- | --- | --- |
| Source | In house | Website* | Website* |
| Instrument | Chromium | GemCode | Chromium |
| Number of Cells | 3,354 | 2,700 | 4,352 |
| Total Number of Reads | 243,335,445 | 185,980,783 | 379,462,522 |
| Total Genes Detected | 18,538 | 16,591 | 19,763 |
| After Preprocessing |  |  |  |
| Number of Cells | 3,199 | 2,591 | 4,300 |
| Mean Counts per Cell | 3,563 | 2,435 | 4,368 |
| Median Genes per Cell | 1,053 | 817 | 1,226 |

\*https://support.10xgenomics.com/single-cell-gene-expression/datasets

Alignment, de-duplication, barcode filtering and gene quantification for all three datasets was handled by the 10X Genomics proprietary software, Cell Ranger. Note that we aligned reads to the hg19 genome annotation. Using the Bioconductor package scater (7), we then removed low quality data from cells with low library size or low number of expressed gene transcripts. We also removed cells with high mitochondrial read proportion as this can indicate apoptosis as such a cell that has an aberrant transcriptome profile in comparison to a living cell. Finally, we also removed any cells that were visible outliers in a plot of number of total features versus log base 10 transformed number of total reads (more detail is given in the Supplementary Information).

### Overview of Single Cell Clustering Methods available in in R applicable to 10X Data

There are many competing clustering methods developed for scRNA-seq data and new methods are constantly emerging in this rapidly developing field. Here, we investigate clustering methods available as R packages. We focused on methods available in the R language, as this is one of the most commonly used programming languages for scRNA-seq data analysis. While we tried over 20 R packages only 11 were able to handle the large size of 10X Genomics scRNA-seq data and had sufficient documentation to allow their application. For some of the R packages the primary focus is not clustering, but the package authors explicitly describe how their packages can be applied to achieve clustering of the scRNA data. We have summarized the main characteristics of the methods in Table 2. Besides methods available as R packages, we also used Cell Ranger, and in particular the graph-based clustering approach available therein. All clustering methods were run using their default settings, including filtering of genes as described in their documentation (for more detail see Supplementary Information). We concede that it is possible that more care in the upstream data handling and selection of parameters could result in different results. However, we explicitly ran these methods in the manner of a non-expert, as this presents the most common user case. Note that for clustering methods that do not automatically choose the number of clusters, we specified 8 clusters.

**Table 2.**
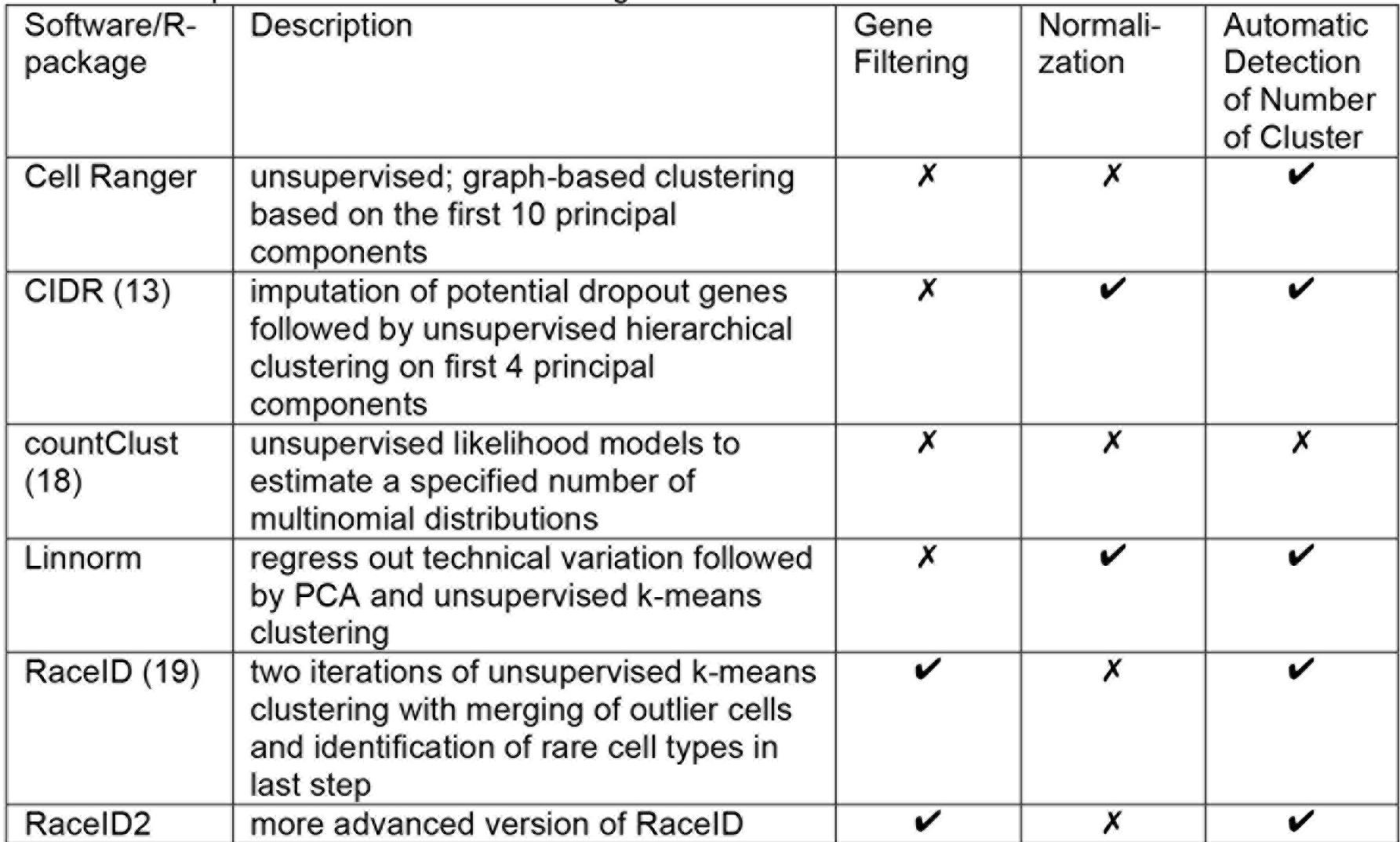
Comparison of different clustering methods

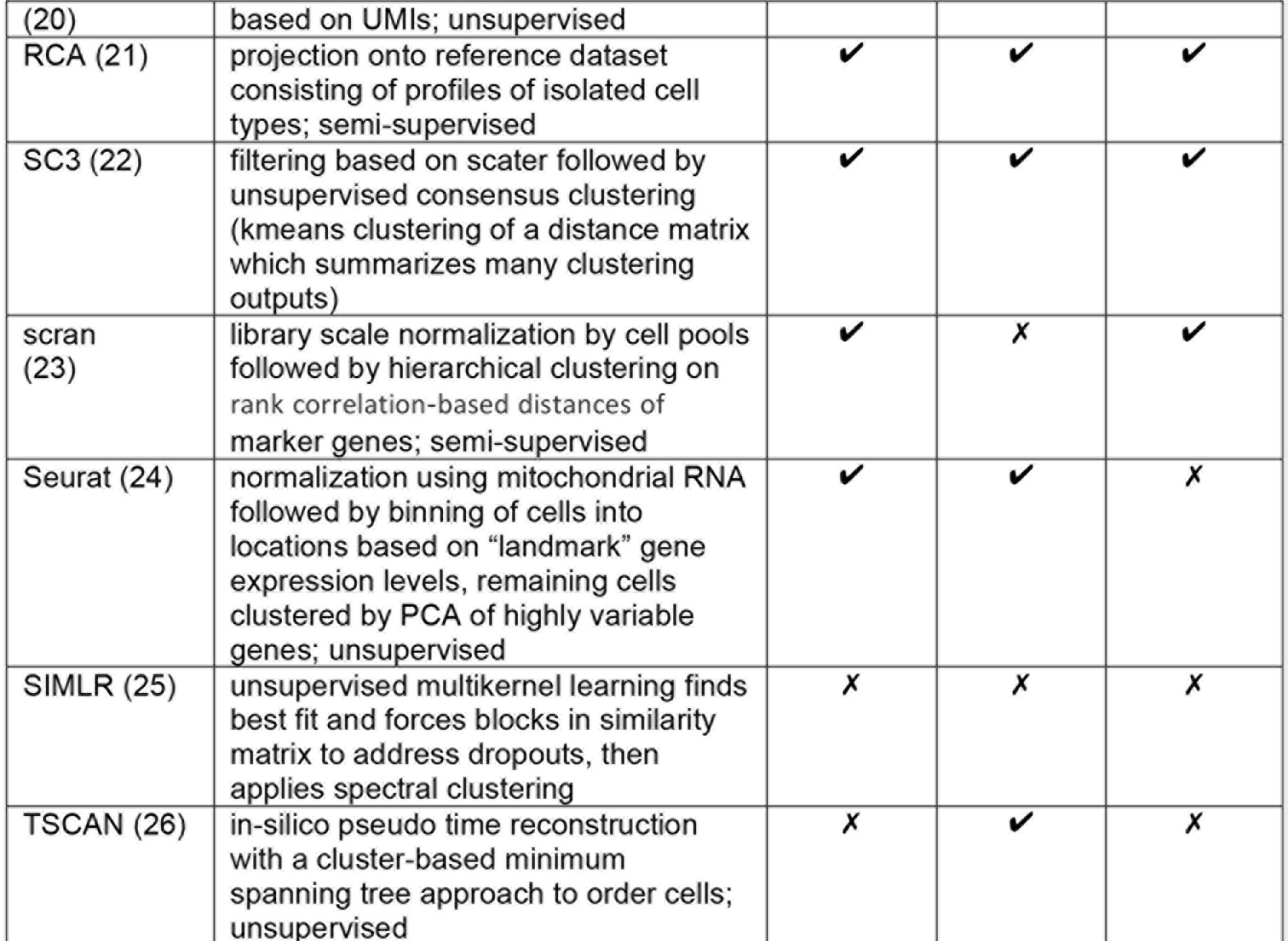

### Evaluating Clustering Solutions in the Absence of Truth

Unlike with some scRNA-seq protocols, with droplet-based protocols, such as 10X Genomics, cell surface markers cannot be used to establish the cell type of each cell. This means that we were forced to compare different clustering methods in the absence of truth. Hence, we investigated the performance of the different methods using three approaches.

#### Examining Robustness of Clustering Strategies

To test the robustness of different clustering methods we pursued a sub-sampling strategy on the largest dataset (10X Chromium). We generated three datasets, each of which included 3000 randomly sampled cells (out of the total of 4,300 that were available after filtering). For every combination of two datasets (three combinations in total) we then investigated for each clustering method separately how often cells contained in all three subsampled datasets were assigned to the same cluster. As cluster labels are meaningless, we used the Adjusted Rand Index (ARI) (8) and the Normalized Mutual Information (NMI) (9), two metrics used routinely in the field of clustering to describe similarity between clustering solutions. Both metrics can take values from 0 to 1, with 0 signifying no overlap between two clustering solutions and 1 signifying complete overlap.

#### Comparing Different Clustering Solutions

The ARI and the NMI can also be used to compare clustering solutions of different methods. We applied both metrics to every pair of clustering methods for all three datasets.

#### Comparing Clustering Solutions to Supervised Labeling

We also compared the clustering solutions by labeling the cell expression profiles of the clusters to average expression profiles from a reference dataset containing 11 isolated cell types for all datasets. While this labeling does not constitute truth it has been found to be consistent with marker-based classification (6). We then use the labels of these clusters derived from this method as a surrogate for the true cell type. In order to assess similarity of any clustering solution with these labels we used two complementary metrics: average homogeneity and average entropy. Average homogeneity can take on values from 0 to 1, with 1 signifying complete agreement. In contrast, average information entropy is positively valued with values closer to 0 signifying more homogeneity in each cluster, i.e. better overlap with the “truth”. Neither metric is ideal, with clustering methods predicting large numbers of clusters favored by the second metric but disfavored by the first, and vice versa for clustering methods predicting few clusters. Hence, we used both metrics in concert to evaluate similarity of each clustering solution with the supervised labeling.

### Assessing Effect of Cell Characteristics on Clustering Solutions

We also investigated what properties of each cell’s data were driving the clustering solutions produced by the different methods. Properties of a cell’s data refer to features such as the number of total reads that included the cell’s barcode, the total number of gene transcripts found for this cell, etc. To this end, we used linear mixed models where cell data properties were predicted using the indicators for cluster membership. We predicted cell data properties and not cluster membership for modeling ease. The adjusted R^2^ of these models were used to assess which properties influenced the clustering solutions from the different clustering methods. Properties investigated included: (i) the total number of detected gene transcripts, (ii) the total read count, and (iii) the percentages of reads aligning to ribosomal protein, mitochondrial and nuclear genes involved in mitochondrial processes (nuclear mitochondrial genes).

## Results

We focused on datasets generated from human PBMCs because they are well studied. They also provide a challenging test framework for the different clustering methods as they contain more than a dozen well-recognized different cell types (10), some of which are closely related. Furthermore, many PBMCs cell types can be isolated, which means useful reference data as well as knowledge on typical PBMC composition exist.

### Clustering Strategies are Robust

The sub-sampling analysis reveals that all clustering methods are reasonably robust to slight changes in input data (see Table 3). RCA unsurprisingly proved particularly robust, because it relies on projection of the cells onto a space defined independently by gene expression of isolated cell types. The least robust method was RaceID2. However, the NMI metric for this method was substantially better than the ARI metric suggesting that the variations between the clustering solutions were small.

**Table 3.** Mean and standard error (SE) of ARI and NMI measuring similarity between clustering solutions produced on 3 different subsampled datasets by different clustering methods. Numbers in bold indicate maximum.

| Software/R-package | ARI ± SE | NMI ± SE |
| --- | --- | --- |
| cellRanger | 0.765 ± 0.130 | 0.839 ± 0.068 |
| CIDR | 0.818 ± 0.066 | 0.827 ± 0.036 |
| countClust | 0.813 ± 0.028 | 0.710 ± 0.042 |
| Linnorm | 0.729 ± 0.081 | 0.800 ± 0.025 |
| RaceID | 0.813 ± 0.028 | 0.710 ± 0.042 |
| RaceID2 | 0.473 ± 0.038 | 0.714 ± 0.016 |
| **RCA** | **0.988 ± 0.005** | **0.982 ± 0.005** |
| SC3 | 0.855 ± 0.009 | 0.886 ± 0.003 |
| Scran | 0.880 ± 0.039 | 0.866 ± 0.029 |
| Seurat | 0.807 ± 0.035 | 0.857 ± 0.010 |
| SIMLR | 0.851 ± 0.019 | 0.866 ± 0.003 |
| TSCAN | 0.706 ± 0.082 | 0.791 ± 0.048 |

### Little Similarity Between Different Clustering Solutions

Comparing different clustering solutions demonstrated that only few a methods resulted in clustering solutions that were similar (see Figure 1 and Figure 2S). For all three datasets, we observed a set of four clustering methods that produced similar results (SIMLR, scran, RCA and Linnorm). A second set, containing eight methods, appeared dissimilar between each other and to the first set. The second set tended to contain methods that estimated higher numbers of clusters. For example, RaceID2, which had little in common with any other clustering approach, estimated more than 100 clusters for all datasets. In order to see whether the differences in number of clusters could explain the dissimilarity of these methods, we set the number of clusters to 8 for all strategies were this was possible (7 out of 12) and re-clustered the cells of the CSL dataset (see Figure 3S). Interestingly, the clustering solutions produced by these methods still remained dissimilar. This indicates that differences between methods are driven not by differences in the estimation of the number of clusters, but rather by differences in methodologies and upstream data handling. We further investigated the effect of upstream data handling, i.e. the application of gene filtering and normalization, by comparing clustering solutions produced by the same methods on differently filtered and normalized versions of the CSL dataset. Again this was not possible for all methods. However, for the methods, which allowed this (6 out of 12), we observed that different upstream handling resulted in markedly different clustering solutions (see Figure 4S).

**Figure 1.**
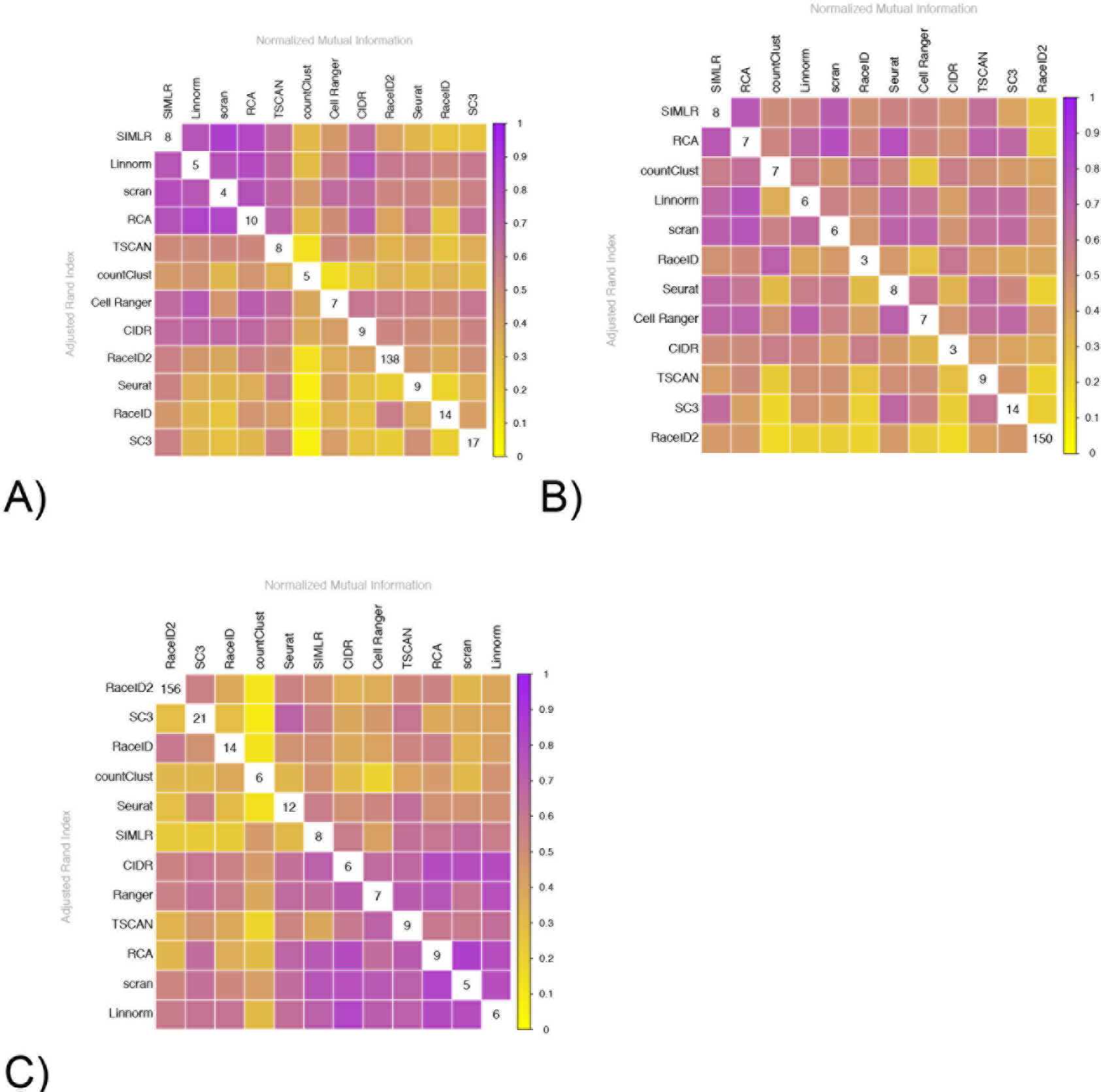
Similarity between clustering solutions of different methods. The lower triangle depicts the ARI of any two methods, while the upper triangle depicts the NMI of any two methods. The numbers on the diagonal give the number of clusters that were estimated. The methods were clustered according to their first principal component of the normalized mutual information. The different panels depict different datasets: A) CSL data B) 10X GemCode data C) 10X Chromium data. For all datasets different methods appear to result in solutions that are different from most other clustering solutions.

### Clustering Strategies Rarely Agree with Supervised Labeling Approach

For all three datasets, the proportions of cells assigned to the 11 cell types by the supervised labeling approach were consistent with the literature (see Table 1S) (11) (12). Furthermore, cell labeling was consistent between the different datasets (compare Figure 2). The datasets generated with the 10X Genomics Chromium system even displayed the same structure for the T-cell subpopulations. The similarity of the datasets is also reflected when assessing the agreement of the clustering solutions and the supervised labeling approach (see Figure 3). For the dataset produced with GemCode, the first 10X microfluidics instrument the labeling was not as consistent, but largely followed the same trend. Generally, none of the clustering methods demonstrated a high homogeneity and low information entropy simultaneously, which would have indicated highest similarity with the supervised labeling solution. Clustering solutions produced by RaceID, SC3, Seurat and TSCAN were the ones most different from the cell assignment of the supervised labeling approach, as these had low homogeneity metrics and high information entropy metrics for all datasets. The failure to replicate the results of the supervised labeling approach shed doubt on the ability of these methods to accurately detect known as well as novel cell populations in 10X Genomics data.

**Figure 2.**
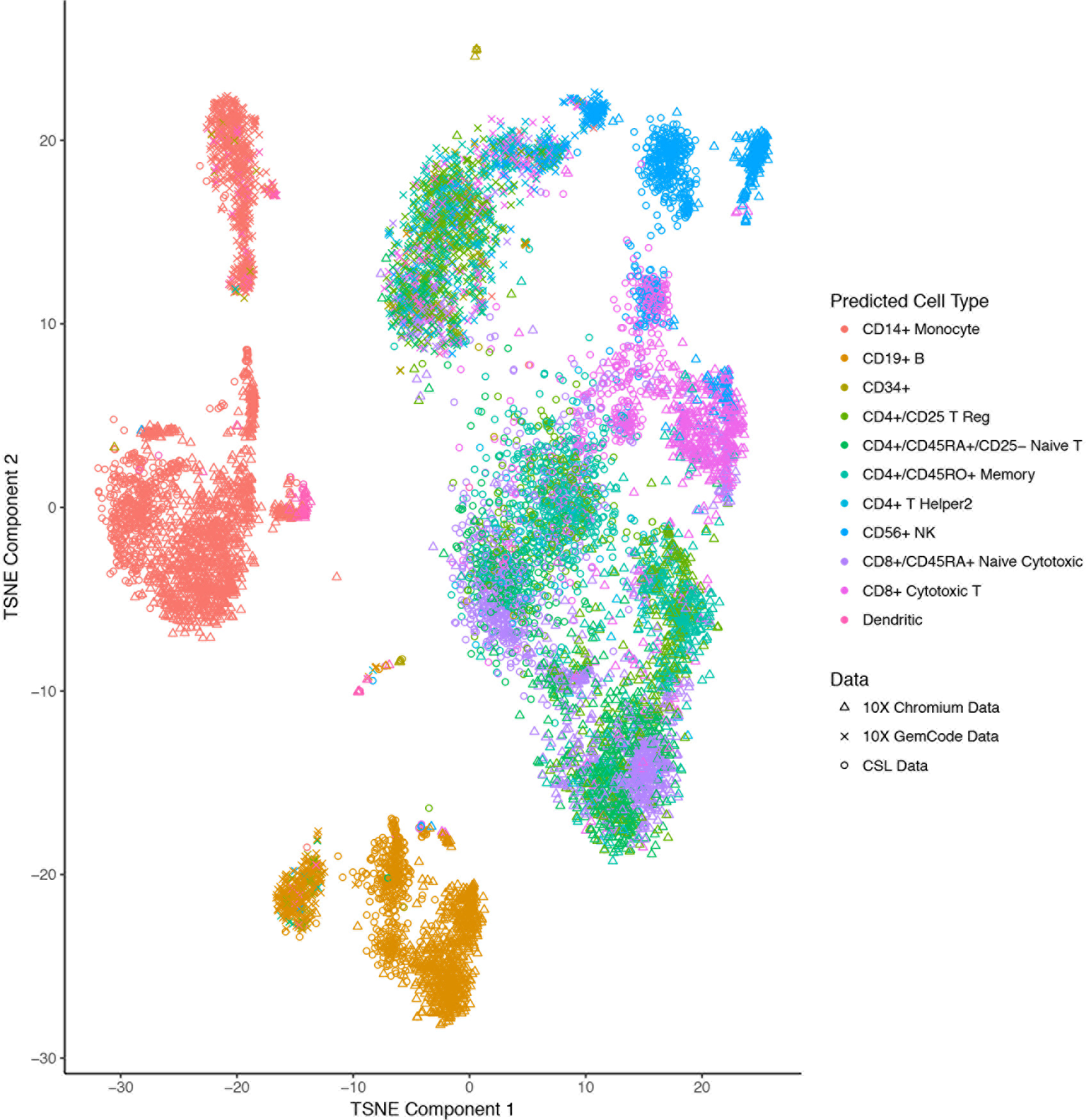
T-SNE representation of cells from all three datasets. The first 500 principal components were estimated for the CSL data. Both 10X Genomics datasets were then projected into this space. Data were then combined and converted to a t-SNE representation (perplexity=100, exaggeration factor=12). (Note that constructing this figure starting in any of the other two datasets qualitatively results in the same figure.) The different datasets are represented by different shapes. Cells were colored with cell type labels from the supervised cell labeling approach described by Zheng et al (6). The figure demonstrates that supervised labeling is consistent between different datasets, because similarly labeled cells from different datasets appear close to each other on the T-SNE representation.

**Figure 3.**
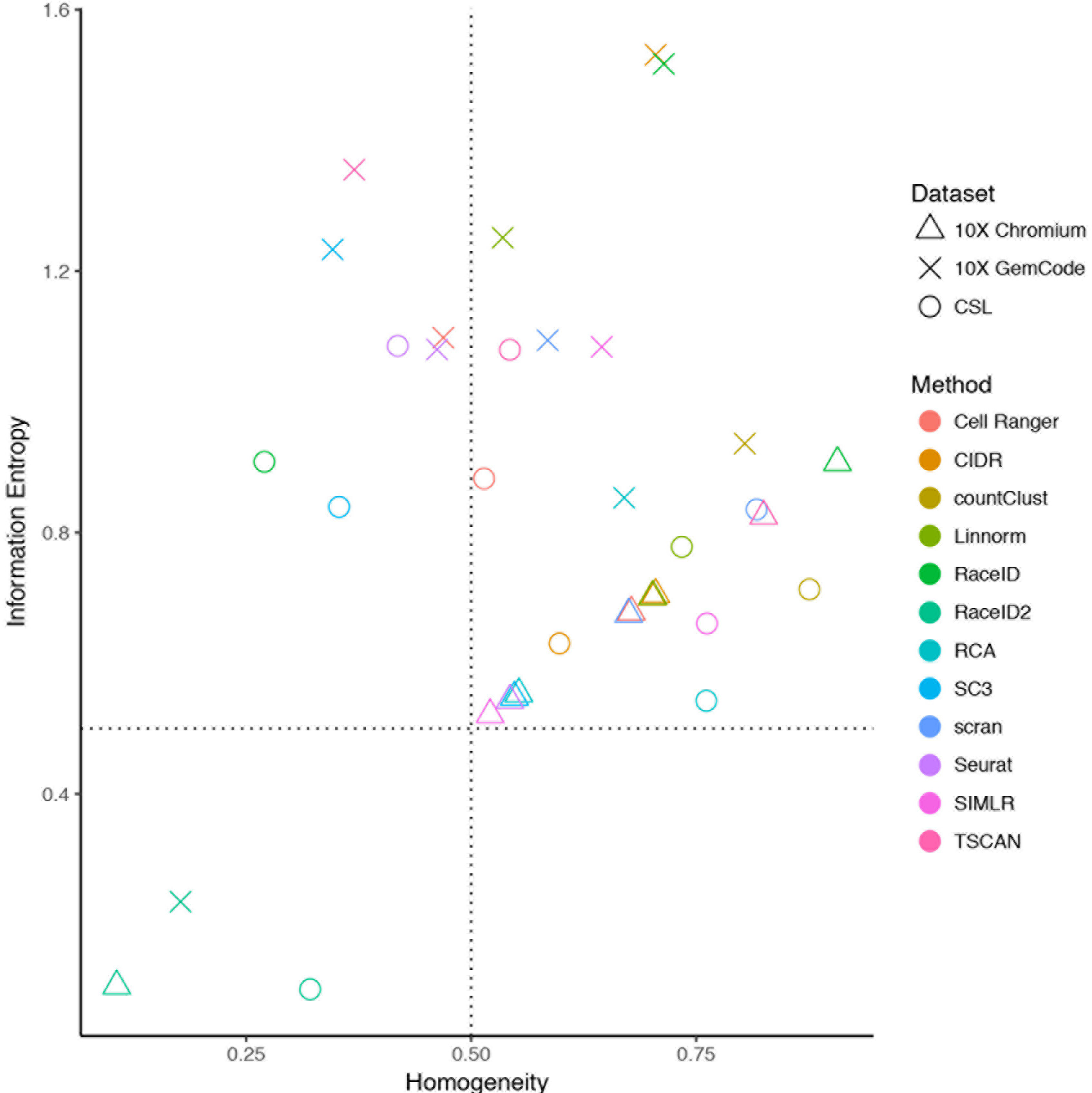
Mean of homogeneity and mean information entropy for each dataset with respect to assignment of cells using supervised cell labeling as described by Zheng et al (6). Different datasets are represented by different shapes, while the color of each point reflects the clustering method. None of the methods achieve high mean homogeneity and low mean information entropy suggesting little overlap with supervised labeling for all methods.

### Clustering Solutions are Driven by Proportion of Ribosomal Protein Genes Sequenced for Each Cell

In all three datasets, variation in the percentage of reads aligning to ribosomal protein genes was the only cell property investigated that strongly predicted by all 12 clustering solutions (compare Figure 4). The one exception is TSCAN, which in addition to being affected by the percentage of reads aligning to ribosomal protein genes was also affected by the total number of detected genes and total number of counts. The method most affected by the percentage of reads mapping to ribosomal protein genes was RaceID2 (R^2^>0.8). This suggests that clustering solutions of all methods foremost reflect differences in the amount of ribosomal protein genes between cells. To which degree this coincides with cell types requires further investigation.

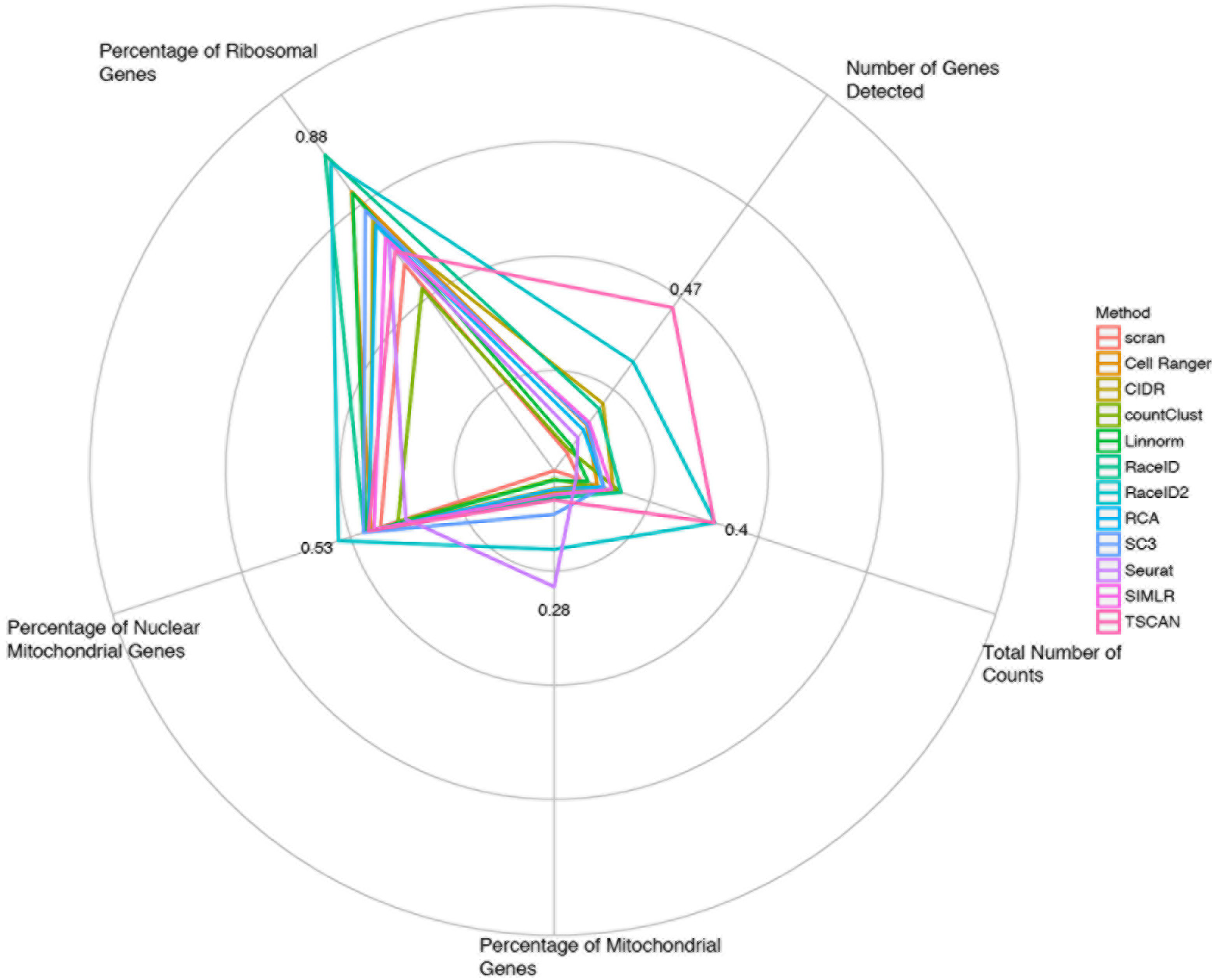

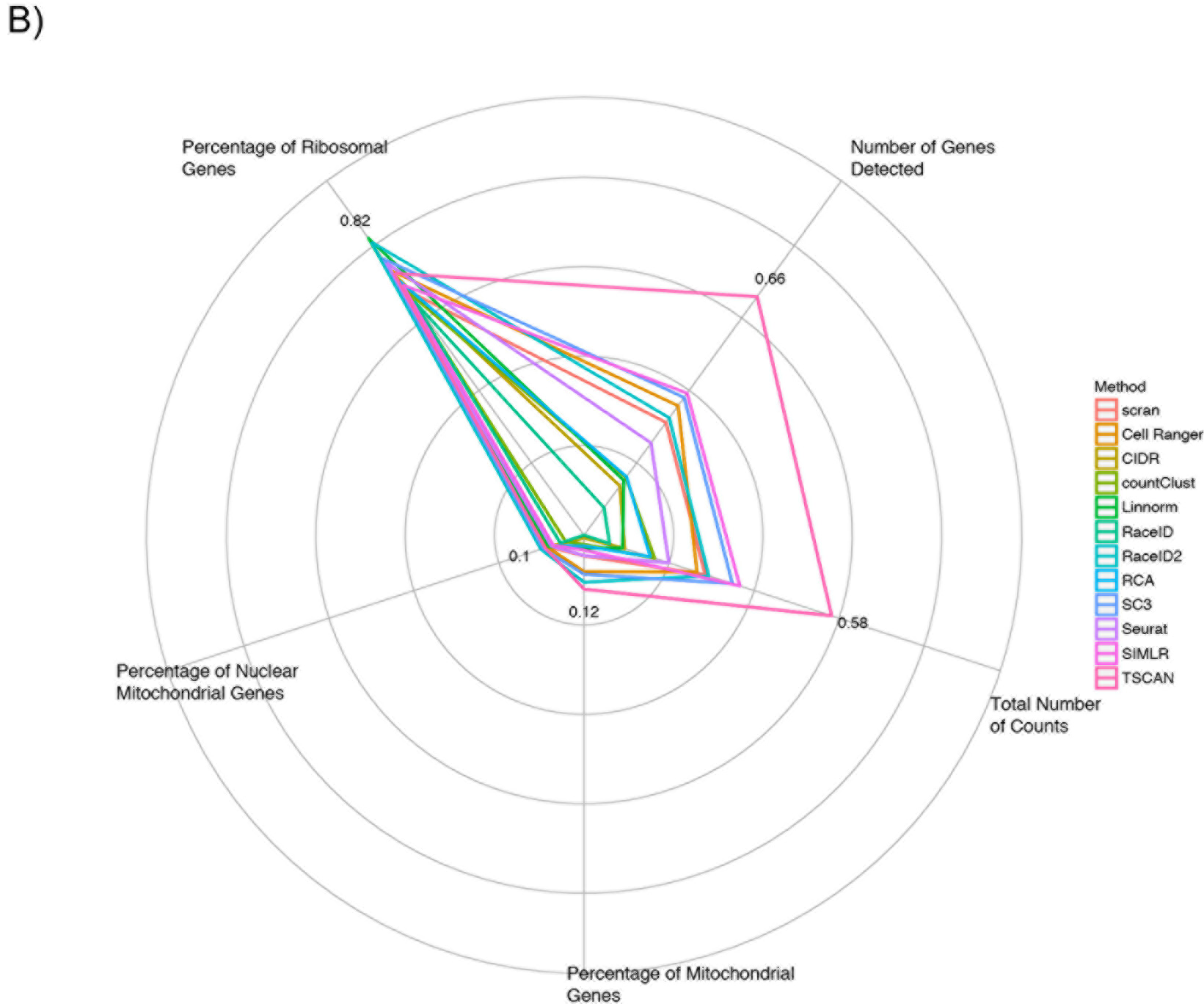

**Figure 4.**
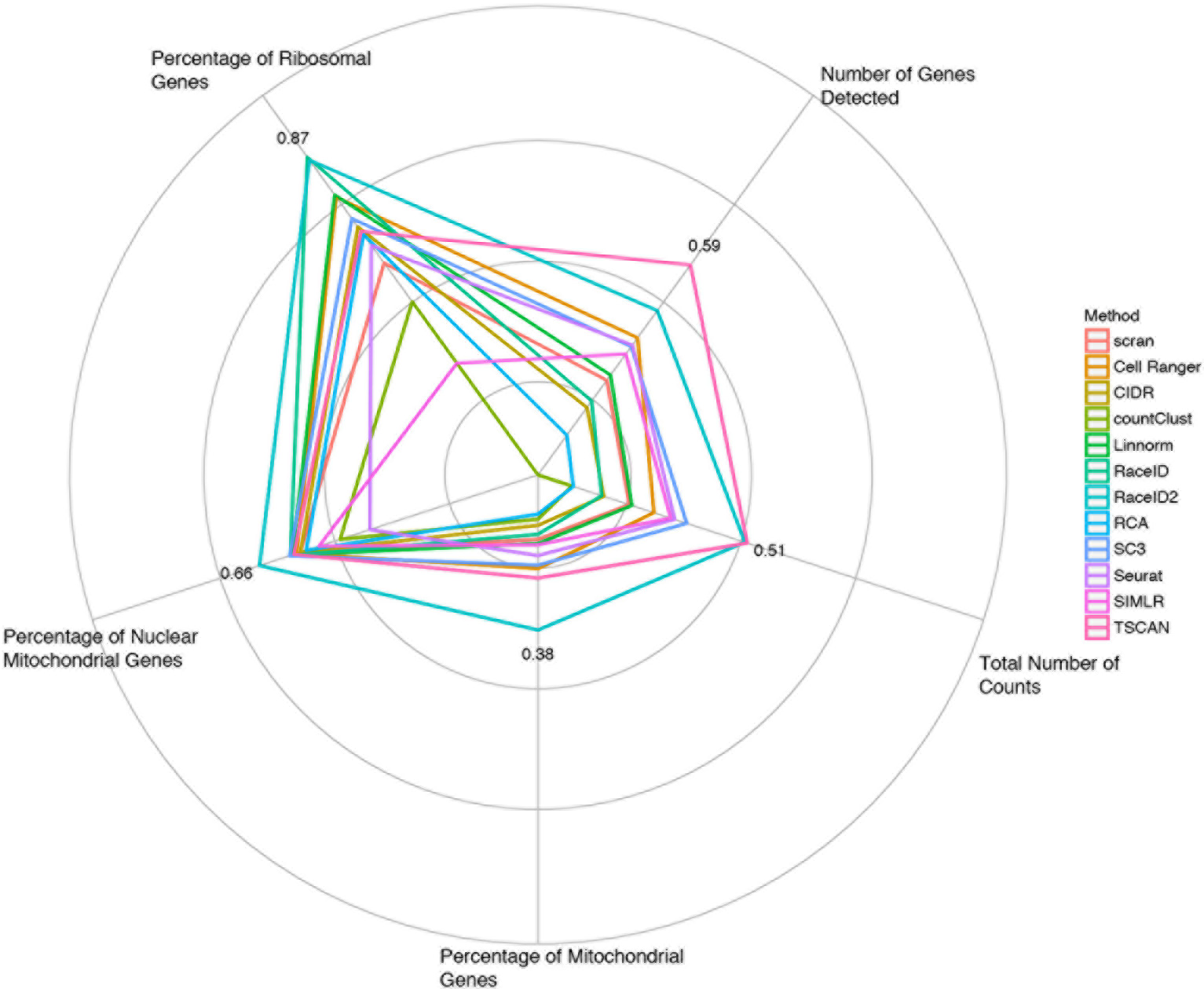
Radial plots describing the effect of 6 cell features on the clustering solutions of different methods. For every method and every feature the adjusted R^2^ of the linear model fitting the feature by the clustering solution is presented. The different panels depict different datasets: A) CSL data All methods are strongly affected by the proportion of ribosomal genes in a cel

Ribosomal protein genes account for around 40% of all reads in all three datasets. They also represent some of the most variable genes. We initially speculated that they dominate the signal. However, when we removed ribosomal protein and mitochondrial genes most methods remained affected by the percentage of reads aligning to ribosomal protien genes (see Table 2S). This suggests that even after removing ribosomal protien and mitochondrial genes, the information about the proportion of reads mapping to these genes is inherently contained in the data. Analogously, removing the ribosomal protien and mitochondrial genes can be thought of as cutting a face from a picture. The hole left by the cut still contains information about the shape, which allows inferences about what has been cut out. Upon removing ribosomal protien and mitochondrial genes the clustering solutions of half of the methods were dramatically changed (see Figure 6S). In the case of CIDR (13) we believe that ribosomal protien genes heavily inform its imputation step resulting in a markedly altered clustering solution. Note that Linnorm failed to cluster the filtered data when run in default mode.

## Discussion

When applied to 10X Genomics scRNA-seq different clustering methods result in different solutions. These can be profoundly different. Thus, it seems likely that different methods cluster this type of data according to different aspects. Our investigations suggest that biological difference between cells, such as cell type or state, and technical variation between cells (as well as combinations of biological differences and technical variation) are all influencing clustering. It is unclear which aspects are captured by which clustering methods. In particular, our results suggest that no method results in clusters that reflect different cell types well, the aspect most sought after.

In order to improve our knowledge about clustering methods more adequate scRNA-seq benchmarking datasets need to be generated. Currently, benchmarking datasets that allow detailed performance analysis of clustering tools only exists for scRNA-seq technologies incorporating FACS. Droplet-based technology, such as 10X Genomics, does not incorporate FACS and consequently cell type labels for each cell are unknown. Traditional mixing experiments can elucidate whether clustering methods are able to accurately estimate proportions of cell types, but they cannot tell us whether any one cell is assigned to the cluster of its type. Nevertheless, in the future CellTagging, where lentiviral transduction is used to introduce unique DNA indexes, might allow generation of droplet-based scRNA-seq data with cell labels (14).

The importance of well-designed benchmarking datasets cannot be overstated. Both microarray quality control studies (15) (16) helped evaluate and develop gene expression analyses for data generated with microarrays. Similarly, the sequencing quality control study (SEQC) informed best practice guidelines and has been used during the development of many innovative methods (17). It is reasonable to anticipate scRNA-seq benchmarking datasets to have similar effects. However, in order for them to be useful to the area of clustering they will need to come with cell labels.

Until benchmarking datasets have been generated, practitioners and consumers of results generated from 10X Genomics scRNA-seq data alike should remain vigilant. The choice of clustering tool for scRNA-seq data generated by the 10X Genomics platform crucially determines interpretation. Hence, at least two clustering tools should be routinely applied to 10X Genomics scRNA-seq data in order to offer more than one subjective interpretation.

## Acknowledgements

We gratefully acknowledge the constructive comments and experimental work of Mark Biondo and Nicolas J. Wilson. We would like to thank the Australian Genome Research Facility for their generous support of this project, including funding. This work was also supported by the Victorian Government’s Operational Infrastructure Support Program and Australian Government NHMRC IRIIS. MB is funded by NHMRC Senior Research Fellowship 110297 and NHMRC ProgramGrant 1054618.

## Data Availability

CSL data: Will be made available upon publication.

10X GemCode data: https://support.10xgenomics.com/single-cell-gene-expression/datasets/1.1.0/pbmc3k

10X Chromium data: https://support.10xgenomics.com/single-cell-gene-expression/datasets/2.0.1/pbmc4k

## Ethics Statement

The IRB of the Bio21 Institue approved the generation of the CSL single cell data (Skin and Cancer Foundation - HREC2012-05-812).

## Programs Used

Cell Ranger: 1.3.1; R: 3.4.0; scater: 1.4.0; cellrangerRkit: 1.1.0; SAVER: 0.1.2; CIDR: 0.1.5; countClust: 0.1.2; Linnorm: 2.0.4; RCA: 1.0.0; SC3: 1.4.2; scran:1.4.2; Seurat: 2.0.1; SIMLR: 1.2.1; TSCAN: 1.14.0

